# Statistical Local Binary Patterns (SLBP): Application to Prostate Cancer Gleason Score Prediction from Whole Slide Pathology Images

**DOI:** 10.1101/503094

**Authors:** Hongming Xu, Tae Hyun Hwang

## Abstract

Computerized whole slide image analysis is important for assisting pathologists in cancer grading and predicting patient clinical outcomes. However, it is challenging to analyze whole slide image (WSI) at cellular level due to its huge size and nuclear variations. For efficient WSI analysis, this paper presents a general texture descriptor, statistical local binary patterns (SLBP), which is applied to prostate cancer Gleason score prediction from WSI. Unlike traditional local binary patterns (LBP) and many its variants, the presented SLBP encodes local texture patterns via analyzing both median and standard deviation over a regional sampling scheme, so that it can capture more micro- and macro-structure information in the image. Experiments on Gleason score prediction have been performed on 317 different patient cases selected from the cancer genome atlas (TCGA) dataset. The presented SLBP descriptor provides over 80% accuracy on two-class (grade ≤7 vs grade ≥8) distinction, which is superior to traditional texture descriptors such as histogram, Haralick and other state-of-art LBP variants.

## 1. INTRODUCTION

With recent advance in digital pathology and computerized algorithms, automatic whole slide image analysis is being actively explored to assist pathologists for cancer grading and predicting patient clinical outcomes. Since pathologists typically diagnose and grade cancer slides by examining cellular structures and its changes, many existing studies attempt to extract feature representations based on cellular analysis. For example, Barker *et al.* [1] proposed a technique for brain tumor sub-typing based on cellular features extracted from local representative tiles of the WSI. Xu *et al.* [2] proposed a technique for automatic diagnosis of skin melanoma from whole slide image analysis, which extracts cellular features from skin epidermis and dermis layers. Since the gigabyte whole slide image usually has millions of cell nuclei with great variations in terms of color, shape and size, extraction of cellular features from WSI is very challenging. Although cellular features can be computed from selected image patches, this may bring the sampling bias for the whole slide image representation.

Because of overwhelming successes in computer vision field, deep learning models have also been applied for whole slide image analysis. Jiménez-del-Toro *et al.* [3] tried several deep learning models including LeNet, AlexNet and GoogLeNet to predict high Gleason scores (i.e., ≥8) from prostate cancer WSI. They evaluated deep learning models on 235 prostate tissue pathology images, and reported about 78% accuracy in distinguishing grades ≥8 and ≤7. Coudray *et al.* [4] reported the success by applying Google inception V3 model on TCGA non-small lung cancer WSI for tumor sub-typing and cancer gene mutation prediction. Although deep learning models are powerful to learn image features, they require exhausted annotations for every training image (e.g., small patches of the WSI). However, the ground truth labels of individual patches are usually unknown, as pathologists typically only assign image-level ground truth label. For example, TCGA prostate cancer patients only have image-level Gleason scores, which were estimated by adding the two dominant tumor grades across the whole tumor region [5]. The lack of annotations for subregions makes it difficult to apply deep learning models for Gleason score prediction on TCGA database.

Since cellular or gland changes can result in texture appearance difference across cancer tissue slides, texture features have also been well-explored for histopathological image analysis. The widely-used textural features are computed from gray level co-occurrence matrix, multi-wavelet transform, fractal analysis and texton maps. Huang *et al.* [6] proposed to apply fractal analysis to describe histological variations within the image, and used it to predict different Gleason grades. Khurd *et al.* [7] proposed to characterize image texture of different tumor Gleason grades by clustering extracted filter responses at each pixel into textons. Almuntashri *et al.* [8] proposed to combine wavelet transform and fractal analysis based features to distinguish Gleason grades 3, 4 and 5. Although these studies reported good performances in prostate cancer grading, they only analyzed selected image patches rather than WSI.

Considering the limitations (e.g., complexity or well annotations) of existing studies in WSI analysis, this paper presents a general texture descriptor, statistical local binary patterns (SLBP), and applies it to predict Gleason scores from prostate cancer WSI. The SLBP is extended from traditional LBP operator, but it can capture more texture information by encoding both regional median and standard deviation values. The rest of paper is organized as follows. Section 2 describes the SLBP descriptor, followed by its application on Gleason score prediction in Section 3. Section 4 provides experimental results, and Section 5 concludes the paper.

## 2. STATISTICAL LOCAL BINARY PATTERNS (SLBP)

The SLBP descriptor characterizes image local structure by statistically encoding the centering patch and those of its surrounding neighboring patches (see Fig. 1). Given an image pixel *g_c_* and its *p* circularly (with radius *r*) and evenly spaced surrounding neighbors *g_n_, n* = 0,1,…, *p* — 1, the SLBP descriptor is computed with 3 components:

> (1) The center pixel representation is:
>
>
> 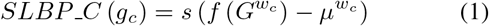
>
>
> where *G^w_c_^* represents a local image patch with a size of *w_c_ × w_c_* and centers at the pixel *g_c_* (see the centering patch with blue borders in Fig. 1). *f* (·) is a statistical function (e.g., median filter or standard deviation function). *μ^w_c_^* denotes the mean of *f (G^w_c_^)* over the whole image with a size of *u × v*, i.e.,
>
>
> 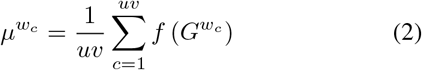
>
>
> and *s* (·) is the sign function.

**Fig. 1:**
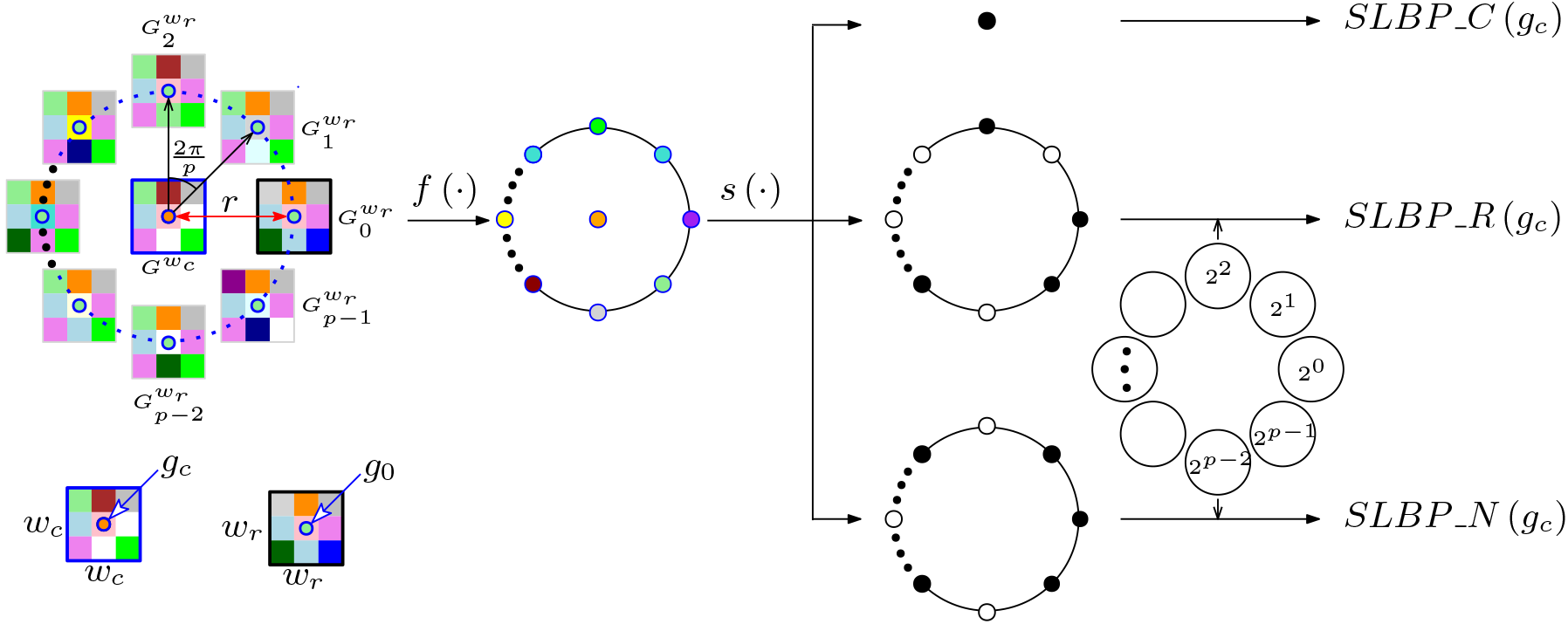
Illustration of statistical local binary pattern (SLBP) descriptor. Note that for illustration the centering patch *G^w_c_^* and the neighboring patch 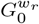 have been shown with blue and black borders. *f* (·) represents statistical function, and *s* (·) represents sign function.

(2)The radial difference representation is:

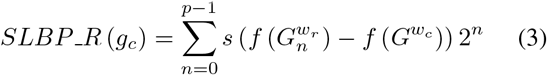

where 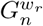 represents a local image patch with a size of *w_r_* × *w_r_* and centers at the neighboring pixel *g_n_* (see surrounding patches in Fig. 1). To ensure the statistical difference in Eq. (3) computed from the same size of patches, we let *w_r_* equal to *w_c_*, i.e.,

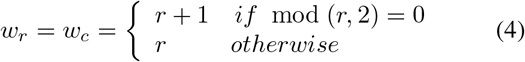

where mod (·) is the modulo function.

(3) The neighbor pixel representation is:

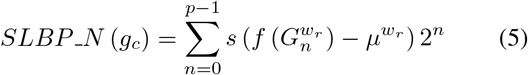

where *μ^Wr^* is the mean over *p* surrounding neighbors 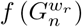, which is computed as:

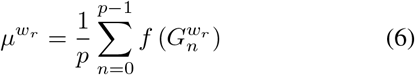

Fig. 1 illustrates pixel encoding by the statistical local binary pattern (SLBP) descriptor. As observed in Fig. 1, × 3 centering and surrounding patches are used for illustration. Since we analyze pixel surrounding patterns over image patches rather than image pixels (like traditional LBP), different statistical functions *f* (·) could be utilized on image patches. In this study, we propose to combine two statistical measures together: median and standard deviation functions. The selection of median function is based on the consideration that median value is robust to image noise and can usually provide the most important intensity information for a certain image patch [9]. The standard deviation function is selected, as standard deviation can usually provide complement information for median value of an image patch.

Note that the *SLBP_C* in Eq. (1) encodes each image pixel value into either 0 or 1. The *SLBP._R* and *SLBP_N* (see Eqs. (3)(5)) encode each image pixel value into the range 0 ~ 2^*p*^ – 1. To reduce the pixel range of *SLBP_R* and *SLBP_N* and make them more robust for texture representation, the rotation invariant uniform *riu*2** encoding scheme [10] is applied, which reduces the values of *SLBP_R* and *SLBP_N* to the range 0 ~ *p* + 1. The joint histograms of feature maps obtained using Eqs. (1)(3)(5) then result in 2(*p* + 2)^2^ dimensional features. The final concatenated features by using both median and standard deviation functions have 4(*p* + 2)^2^ dimensions per image.

Like most other LBP variants, the SLBP descriptor can achieve any spatial resolution and angular resolution analysis by adjusting (*r, p*) values. Fig. 2 illustrates the sampling scheme of our multi-scale SLBP descriptors. In Figs. 2(a)-(d) the spatial scale of SLBP descriptor is increased from *r*_1_ to *r*_4_. Since image texture can be interpreted as textons with different scales, multi-scale analysis by using different (*r, p*) values can help in capturing both micro- and macro-structure information in the image and usually improve classification performance. The features obtained from multi-scale analysis are concatenated together to represent image texture, which results in 4*m*(*p* + 2)^2^ dimensional features, where *m* (e.g., *m* = 4 in this study) indicates the number of scales applied.

**Fig. 2:**
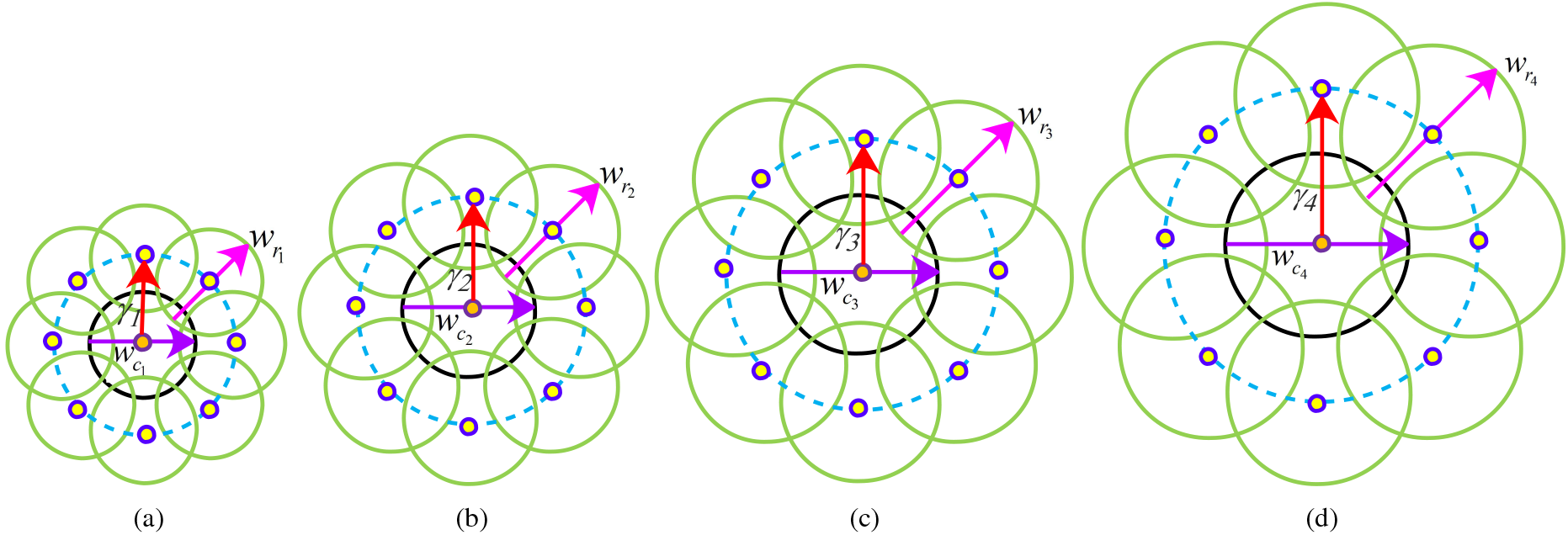
Illustration of sampling schemes for multiscale SLBP descriptor. (a) (*r*_1_, *p*). (b) (*r*_2_, *p*). (c) (*r*_3_, *p*). (d) (*r*_4_, *p*). Note that black and green circles represent centering and neighboring patches, respectively. In all scales, *w_r_* = *w_c_* (e.g., *w_r_1__ = w_c_1__*) and *p* = 8.

## 3. APPLICATION TO GLEASON SCORE PREDICTION

In this section, we apply SLBP descriptor on WSI to predict prostate cancer Gleason scores. Since the WSI with high resolution (e.g., 40×) has a very large size, it is challenging to be directly processed with computerized algorithms. For efficient processing, we employ a multi-resolution analysis strategy to compute features and predict Gleason scores from the WSI. The brief steps are as follows.

(1) Otsu’s multi-thresholding method is performed on the WSI with 2.5 × magnification, which segments image pixels into one of three classes: background, stroma and nuclei (see Fig. 3 (b)). The image within the bounding box of prostate tissue regions is then divided into many non-overlapping tiles. The tiles containing high density of nuclei pixels (i.e., over 25% of tile pixels) are selected. As shown in Fig 3(d), the blue squares highlight selected image tiles which mainly belong to tumor regions of the slide image.

**Fig. 3:**
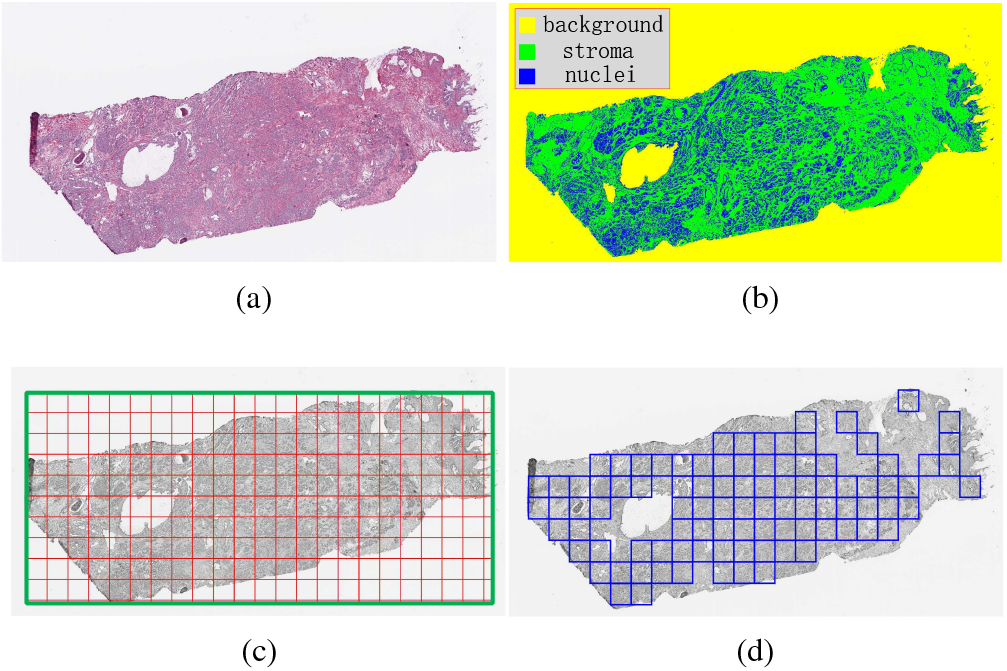
Illustration of tumor tile selection. (a) A H&E stained prostate WSI. (c) Image segmentation, where yellow, green and blue pixels represent image background, tissue stroma and nuclei regions, respectively. (c) Prostate tissue bounding box and divided tiles (128 × 128 pixels). (d) Selected image tiles.

(2) Following the recommendation in [5], all selected image tiles (red channels) with 5× magnification are used to extract texture features using multi-scale SLBP descriptor (see Fig. 2). In this study, the SLBP parameters are set as: *r* = {2,4, 6, 8} and *p* = 8 for 4 spatial scales. The features computed across selected image tiles are then averaged as feature vector representation for the WSI. Each WSI finally corresponds to a 1600 dimensional feature vector.

(3) Since WSI has a high dimension of feature representation, the principal component analysis (PCA) is first applied to reduce feature dimensions to against over-fitting problem. The SVM classifier with a linear kernel is then applied to prostate cancer Gleason score prediction.

## 4 EXPERIMENTAL RESULTS

In this section, we illustrate the results of prostate cancer Gleason score prediction using SLBP descriptor and its comparisons. For this study, a total of 317 whole slide prostate cancer images were obtained from TCGA, which were stained with hematoxylin and eosin (H&E). Each pathology slide had been digitized at multiple resolutions ranging from 2.5 to 40 ×, with all images having a maximum scanning resolution scan at least 20 ×. The ground truths of Gleason scores are provided by TCGA on patient records. Among 317 patient slides, 37 patients are diagnosed as low risk with Gleason score 6, 141 patients are diagnosed as intermediate risk with Gleason score 7 and 139 patients are diagnosed as high risk with Gleason score ≥ 8. To ensure roughly balanced classification, this study evaluates two-class distinction: Gleason scores ≤ 7 (negative) versus ≥ 8 (positive).

To verify the efficacy of presented SLBP descriptor, it has been compared to several widely-used texture descriptors for pathology image analysis. These compared texture descriptors include Fractal Analysis (FA) [6], Histogram, Haralick and Gabor filter (HHG) [11], LBP [10], completed local binary pattern (CLBP) [12] and median robust extended local binary pattern (MRELBP) [9]. For comparison, all texture descriptors combined with SVM classifier with a linear kernel are used for prostate cancer Gleason score prediction. Since the FA [6] only has 8 dimensional features, PCA has not applied on it. However, PCA has been applied on all other texture descriptors before classification. In this study, the feature dimension is empirically reduced to 40 after PCA transform, as it provides the best performance than other empirically reduced feature dimensions (e.g., 30 or 50) for all texture descriptors.

The evaluation of two-class distinction is performed by using recall (*Rec*), specificity (*Spe*) and accuracy (*Acc*) [13]. In addition, we also compute the area under the receiver operating characteristic (ROC) curve to evaluate two-class distinction performance, which is denoted by *Auc*. We perform 5-fold cross validation and repeat it 50 times for all comparisons. The average values of different evaluation criteria are provided.

Table 1 lists classification results obtained by different texture descriptors. As observed in Table 1, the FA [6] provides the poorest performance in terms of *Acc* and *Auc* values among all techniques. The HHG [11] and LBP [10] provide better performances than the FA descriptor, but poorer performances than two state-of-the-art LBP variants, CLBP [12] and MRELBP [9]. The presented SLBP provides steadily improvements over all compared existing texture descriptors, which provides over 80% accuracy and 0.87 *Auc* value. The improvements for the presented SLBP descriptor are mainly because both median and standard deviation of regional patches are used for texture encoding, while existing LBP variants such as CLBP and MRELBP only encode pixel intensities. Fig. 4 shows the ROC curve for two-class distinctions (≤ 7 vs ≥ 8) by using different texture descriptors. As seen in Fig. 4, the SLBP descriptor provides larger Auc values than all other texture descriptors in predicting high risk prostate cancer patients.

**Table 1:**
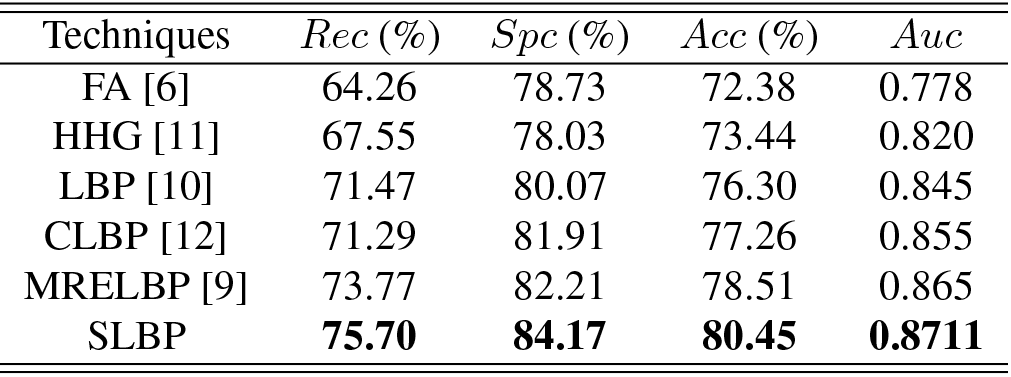
Comparisons on Gleason score prediction

**Fig. 4:**
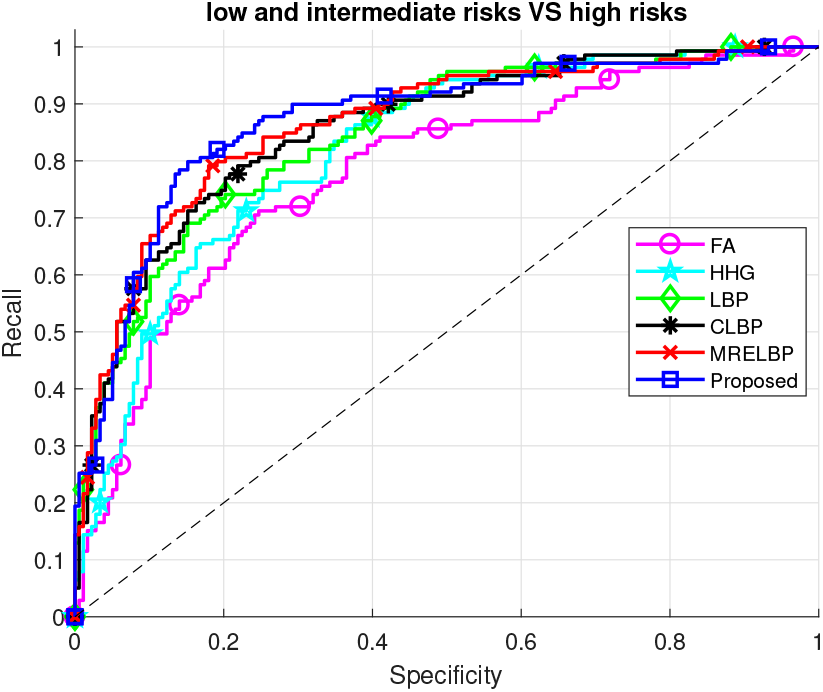
ROC curves for different texture descriptors.

## 5. CONCLUSION

In this paper, we present a general texture descriptor, statistical local binary patterns (SLBP), and applies it to predict prostate cancer Gleason scores from H&E stained whole slide pathology images. The SLBP descriptor encodes local texture patterns by using both median and standard deviation values in a regional sampling scheme, which can capture more micro- and macro-structure information in the image. Experiments performed on a TCGA cohort with 317 different patient pathology images for two-class (≤ 7 vs ≥ 8) distinction indicate the superior performance of SLBP descriptor over state-of-the-art LBP variants and other widely-used texture descriptors such as Haralick and Gabor filters.

## REFERENCES

[1] Jocelyn Barker, Assaf Hoogi, Adrien Depeursinge, and Daniel L Rubin, “Automated classification of brain tumor type in whole-slide digital pathology images using local representative tiles,” Medical image analysis, vol. 30, pp. 60–71, 2016.

[2] Hongming Xu, Cheng Lu, Richard Berendt, Naresh Jha, and Mrinal Mandal, “Automated analysis and classification of melanocytic tumor on skin whole slide images,” Computerized Medical Imaging and Graphics, vol. 66, pp. 124–134, 2018.

[3] Oscar Jiménez del Toro, Manfredo Atzori, Sebastian Otalora, Mats Andersson, Kristian Eurén, Martin Hed-lund, Peter Rönnquist, and Henning Müller, “Convolutional neural networks for an automatic classification of prostate tissue slides with high-grade gleason score,” in Medical Imaging 2017: Digital Pathology. International Society for Optics and Photonics, 2017, vol. 10140, p. 1014000.

[4] Nicolas Coudray, Paolo Santiago Ocampo, Theodore Sakellaropoulos, Navneet Narula, Matija Snuderl, David Fenyö, Andre L Moreira, Narges Razavian, and Aristotelis Tsirigos, “Classification and mutation prediction from non-small cell lung cancer histopathology images using deep learning,” Nature medicine, p. 1, 2018.

[5] Jennifer Gordetsky and Jonathan Epstein, “Grading of prostatic adenocarcinoma: current state and prognostic implications,” Diagnostic pathology, vol. 11, no. 1, pp. 25, 2016.

[6] Po-Whei Huang and Cheng-Hsiung Lee, “Automatic classification for pathological prostate images based on fractal analysis,” IEEE transactions on medical imaging, vol. 28, no. 7, pp. 1037–1050, 2009.

[7] Parmeshwar Khurd, Claus Bahlmann, Peter Maday, Ali Kamen, Summer Gibbs-Strauss, Elizabeth M Genega, and John V Frangioni, “Computer-aided gleason grading of prostate cancer histopathological images using texton forests,” in, 2010 IEEE International Symposium on Biomedical Imaging: From Nano to Macro. IEEE, 2010, pp. 636–639.

[8] Ali Almuntashri, Sos Agaian, Ian Thompson, Danny Rabah, Osman Zin Al-Abdin, and Marlo Nicolas, “Gleason grade-based automatic classification of prostate cancer pathological images,” in 2011 IEEE International Conference on Systems, Man, and Cybernetics (SMC). IEEE, 2011, pp. 2696–2701.

[9] Li Liu, Paul Fieguth, Matti Pietikäinen, and Songyang Lao, “Median robust extended local binary pattern for texture classification,” in 2015 IEEE International Conference on Image Processing (ICIP). IEEE, 2015, pp. 2319–2323.

[10] Timo Ojala, Matti Pietikainen, and Topi Maenpaa, “Multiresolution gray-scale and rotation invariant texture classification with local binary patterns,” IEEE Transactions on pattern analysis and machine intelligence, vol. 24, no. 7, pp. 971–987, 2002.

[11] Jakob Nikolas Kather, Cleo-Aron Weis, Francesco Bianconi, Susanne M Melchers, Lothar R Schad, Timo Gaiser, Alexander Marx, and Frank Gerrit Zöllner, “Multi-class texture analysis in colorectal cancer histology,” Scientific reports, vol. 6, pp. 27988, 2016.

[12] Zhenhua Guo, Lei Zhang, and David Zhang, “A completed modeling of local binary pattern operator for texture classification,” IEEE Transactions on Image Processing, vol. 19, no. 6, pp. 1657–1663, 2010.

[13] Hongming Xu and Mrinal Mandal, “Epidermis segmentation in skin histopathological images based on thickness measurement and k-means algorithm,” EURASIP Journal on Image and Video Processing, vol. 2015, no. 1,pp. 18,2015.

